# Mapping of dI3 neuron sensorimotor circuits across the cervical and lumbar spinal cord

**DOI:** 10.1101/2024.11.17.624039

**Authors:** Shahriar Nasiri, Alex M. Laliberte, Tuan V. Bui

**Affiliations:** Department of Biology, University of Ottawa, Ottawa, ON, Canada; Brain and Mind Research Institute, University of Ottawa, Ottawa, ON, Canada

## Abstract

From the fine control of hand movements to the dynamic corrective adjustments during locomotion, spinal circuits integrate descending supraspinal and sensory inputs to modulate diverse motor functions. The integration of such a wide range of signals across the spinal cord is primarily mediated by propriospinal interneurons. In this study, we investigate the connectivity of a population of propriospinal interneurons marked by the expression of *Isl1*, called dI3 neurons. These dI3s integrate supraspinal and sensory signals to facilitate many important functions, such as hand grasp, locomotion, and motor recovery after spinal cord injury; however, we have a limited understanding of how subpopulations of dI3s modulate network activity across the spinal cord to contribute to these behaviors. Their functional connectivity to motor circuits across the cervical and lumbar spinal cord was assessed through optogenetic activation of dI3s localized in different spinal segments. Our data demonstrates that cervical and lumbar dI3 subpopulations can form local, commissural, intersegmental, and long propriospinal pathways. Furthermore, dI3 subpopulations can be tonically stimulated to elicit locomotor activity. These extensive projection patterns of dI3s across the cervical and lumbar spinal cord suggest that dI3 subpopulations can modulate the activity of multiple motor networks within their respective spinal cord segments or across distant forelimb and hindlimb segments to facilitate a wide variety of motor functions.

## Introduction

Balanced locomotion requires the coordination of many muscles across the body and limbs. While the brain forms descending pathways to the spinal cord crucial for planning and initiating locomotor activity, many functions involved in generating and modulating locomotor output are performed locally by networks within the spinal cord (Kiehn, 2006). These locomotor networks perform distinct functions, such as rhythm and pattern generation, and facilitate the appropriate timing of muscle activity to maintain interlimb and flexor-extensor coordination at various speeds (Bellardita & Kiehn, 2015; Kiehn, 2011, 2016; Krouchev et al., 2006).

Although the spinal locomotor network can produce patterned activity that resembles locomotion without sensory feedback (Marder & Bucher, 2001), adaptable and balanced movements in a dynamic environment require the integration of proprioceptive and mechanoreceptive inputs by sensorimotor circuits (Rossignol et al., 2006). While reflex pathways can facilitate some of these sensory-mediated motor responses by directly activating motoneurons, more complex behaviours that require modulation of motor activity that engages multiple muscles acting across several joints are performed by propriospinal interneurons (PINs), which receive multiple sensory feedback modalities and project to multiple spinal segments (Akay et al., 2014; Laliberte et al., 2019). These propriospinal interneurons are made up of several molecularly defined populations, each with their specific connectivity patterns and role in locomotion (Bui et al., 2016; Danner et al., 2019; Dougherty et al., 2013; Dyck et al., 2012; Koch et al., 2017; Russ et al., 2021; H. Zhang et al., 2022; J. Zhang et al., 2014; Y. Zhang et al., 2008).

Propriospinal interneurons can form local intrasegmental circuits though they play a key role in relaying supraspinal and sensory commands within and across spinal cord cervical and lumbar enlargements (Laliberte et al., 2019; Ueno et al., 2018). Within the enlargements, short PINs can form connections 1-6 segments bidirectionally to convey supraspinal commands, relay local sensory feedback, and propagate locomotor activity across adjacent spinal segments (Laliberte et al., 2019). Some short PINs primarily project ipsilaterally, functioning to coordinate timing of intralimb muscles (Britz et al., 2015; Bui et al., 2016; Dougherty et al., 2013; Laliberte et al., 2019; J. Zhang et al., 2014), while other short PINs form commissural pathways to coordinate the left-right limb pairs (Danner et al., 2019; Dyck et al., 2012; Lanuza et al., 2004; Rabe et al., 2009; Talpalar et al., 2013). Conversely, long PINs form cervicolumbar and lumbrocervical connections to relay supraspinal, sensory, and motor commands between spinal enlargements to coordinate specific hindlimb and forelimb muscles (Brockett et al., 2013; Flynn et al., 2017; Pocratsky et al., 2020; Ruder et al., 2016; Shepard et al., 2023).

The dI3 neurons are one such population of PINs that facilitates some of these sensorimotor roles (Bui et al., 2013, 2016). These neurons are a dorsally-derived class of glutamatergic spinal interneurons marked by the expression of *Isl1* (Bui et al., 2013). These dI3s receive inputs from the primary motor cortex, integrate sensory feedback from cutaneous low-threshold mechanoreceptors and proprioceptors, form premotor circuits with motoneurons, and activate locomotor networks (Bui et al., 2013, 2016; Ueno et al., 2018). dI3s have been implicated in various motor tasks requiring sensory integration, such as palm grasping, locomotion, and locomotor recovery after spinal cord injury (Bui et al., 2013, 2016). These diverse roles of dI3s suggest their involvement in several propriospinal pathways that relay supraspinal, sensory, and locomotor commands across several spinal segments and potentially, across spinal enlargements (Bui et al., 2013, 2016; Goetz et al., 2015; Pivetta et al., 2014; Stepien et al., 2010; Ueno et al., 2018).

While the general connectivity of dI3s have been characterized and certain roles were identified (Bui et al., 2013, 2016), an understanding of how dI3s across the spinal cord convey brain and sensory commands within and across spinal enlargements is limited. This study aims to characterize the functional connectivity patterns of these dI3s located across the cervical and lumbar spinal cord to motoneurons and spinal locomotor networks.

## Results

To map the connectivity of dI3s throughout cervical and lumbar segments, we developed a transgenic mouse line, *Isl1^Cre^*^+/-^; *Slc17a6^FlpO^*^+/+^; *Ai80^(RCFL-CatCh)-D^* (dI3-driver-CatCh), that genetically expresses a channelrhodopsin, called CatCh (Kleinlogel et al., 2011), in Isl1^+^/Vglut2^+^ (encoded by the gene *Slc17a6*) cells (Figure 1A). In addition to dI3s, this genetic approach leads to the expression of CatCh in Vglut2^+^ nociceptive afferents (Figure 1B). To minimize the activation of these afferents, we performed optogenetic experiments in isolated neonatal P1-P7 spinal cords with dorsal hemisections at lumbar and cervical enlargements (Figure 1B & C). Three suction electrodes were applied simultaneously to selected lumbar and cervical ventral roots with the spinal cord positioned ventrally upward; however, all ventral roots from C4-C8 and L1-L5 were sampled across n = 11 mice (Figure 1C). Blue-light illumination was applied to spinal segments that were either ipsilateral or contralateral to the recorded ventral roots (Figure 1C). To avoid recruiting polysynaptic networks through continuous dI3 activation, we applied short 10 ms blue-light pulses every 4 seconds for 10 sweeps.

**Figure 1.**
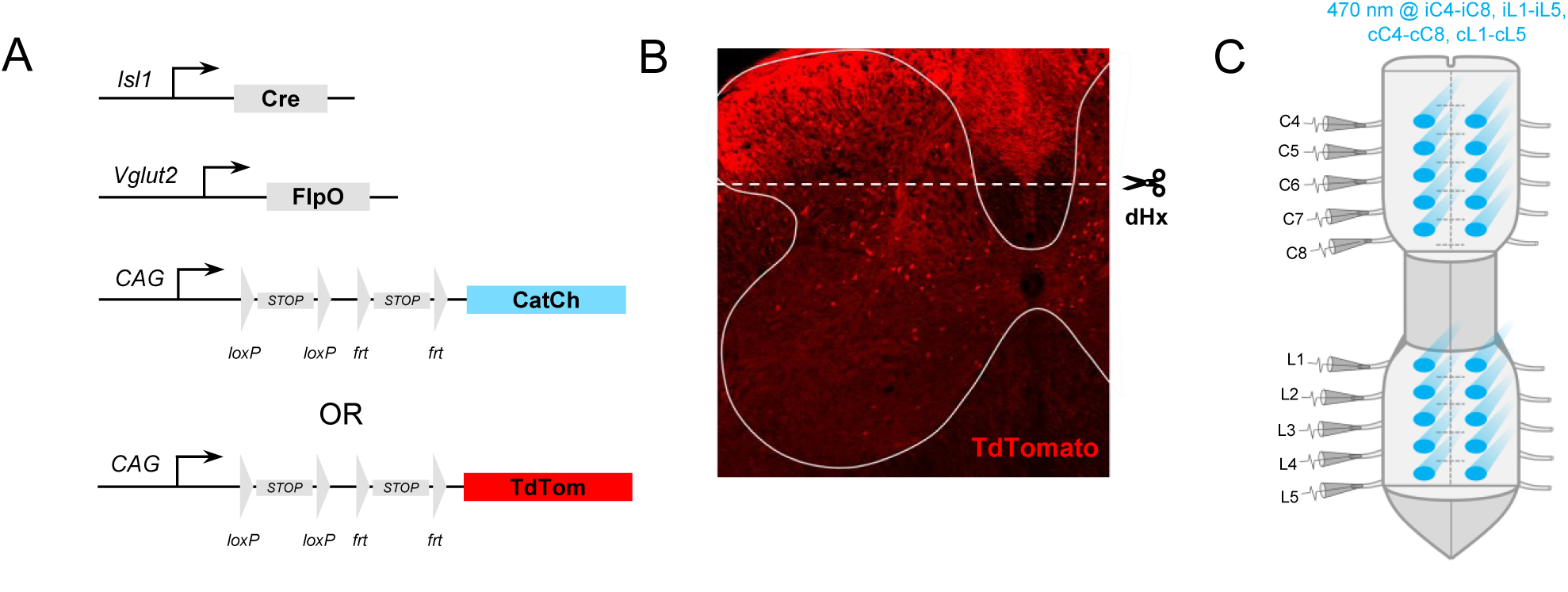
Strategy for optogenetic stimulation of segmental dI3 subpopulations. A. Diagram of genetic strategy for generating *Isl1^Cre^*^+/-^; *Slc17a6^FlpO^*^+/+^; *Ai80^(RCFL-CatCh)-D^* and *Isl1^Cre^*^+/-^; *Slc17a6^FlpO^*^+/+^; *Ai65^(RCFL-tdT)-D^* mice. *Isl1^Cre^*^+/-^ mice were crossed with *Slc17a6^FlpO^*^+/+^ mice to drive the expression of Cre and FlpO recombinase in dI3 neurons and Vglut2^+^ nociceptive afferents, forming dI3-driver mice. For optogenetic experiments, *Isl1^Cre^*^+/-^; *Slc17a6^FlpO^*^+/+^ (dI3-driver) mice were crossed with *Ai80^(RCFL-CatCh)-D^* (CatCh) mice containing a *frt*-flanked STOP cassette and a *loxP*-flanked STOP cassette upstream of the CatCh/EYFP fluorescent protein, resulting in *Isl1^Cre^*^+/-^; *Slc17a6^FlpO^*^+/+^; *Ai80^(RCFL-CatCh)-D^* (dI3-driver-CatCh) mice. To visualize the dI3 neurons and Vglut2^+^ afferents, dI3-driver mice were crossed with *Ai65^(RCFL-tdT)-D^* (TdTomato) mice containing a *frt*-flanked STOP cassette and a *loxP*-flanked STOP cassette upstream of the TdTomato fluorescent protein, resulting in *Isl1^Cre^*^+/-^; *Slc17a6^FlpO^*^+/+^; *Ai65^(RCFL-tdT)-D^* (dI3-driver-TdTomato) mice. B. Representative image of TdTomato (red) labelled dI3 neurons and Vglut2^+^ afferents in dI3-driver-TdTomato mice. To avoid activation of these afferents in dI3-driver-CatCh mice, a dorsal hemisection was performed at the indicated level. C. Diagram of the *ex vivo* spinal cord preparation. The spinal cord was isolated in neonatal P1-P7 dI3-driver-CatCh mice, and a dorsal hemisection was performed at the cervical (C1-C8) and lumbar (L1-L5) segments to remove Vglut2-CatCh^+^ afferents. Three suction electrodes were simultaneously applied to various ventral roots to record from C4-C8 and L1-L5 segments, and blue light was applied to ipsilateral segments. Within the same animals, blue light was also independently applied to contralateral segments from C4-C8 and L1-L5.

### Cervical motoneurons receive input from cervical and lumbar dI3s

To explore which dI3 subpopulations across the spinal cord influence the motor response in the cervical cord, we independently activated ipsilateral C4-C8 and L1-L5 segments while recording from C4-C8 ventral roots (Figure 2B). We took the average of the 10 sweeps of each recording and identified subpopulations of dI3s across the cervical and lumbar enlargement with distinct activation patterns of cervical motoneuron pools (Figure 2A).

**Figure 2.**
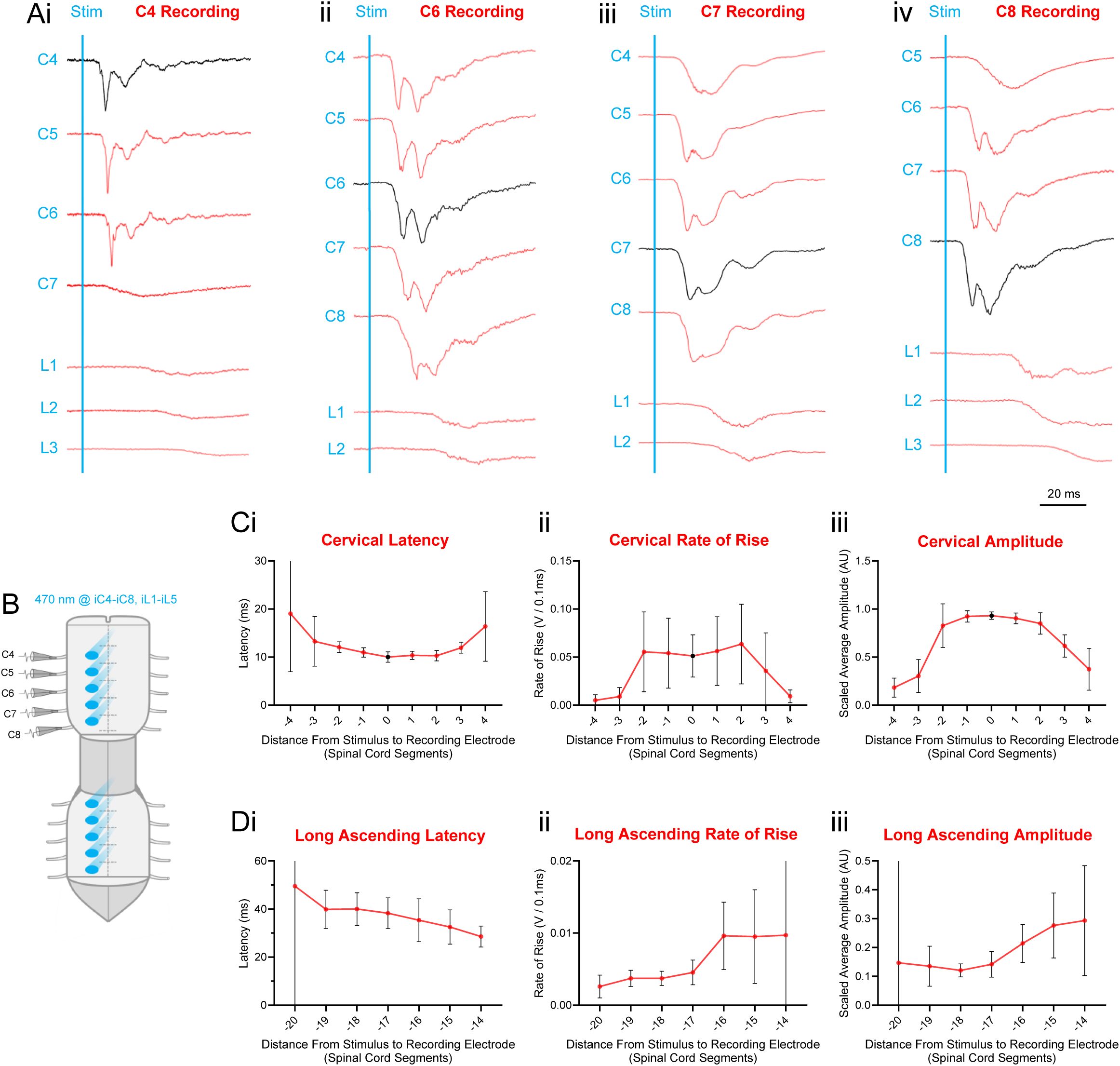
Cervical motoneurons receive input from cervical and lumbar dI3 neurons. A. Averaged traces of cervical ventral root recordings in response to 10 ms optogenetic stimulations of ipsilateral dI3 neurons located from segments C4-C8 and L1-L5. Recordings of optogenetic stimulation of intrasegmental dI3s (black trace), rostral dI3s (red traces above), and caudal dI3s (red traces below) relative to the recorded cervical ventral root are shown. Short-latency, fast-rising, high-amplitude motor responses were observed in: (i) C4 with stimulation of C4-C6 (intrasegmental and up to 2 caudal segments), (ii) C6 with stimulation of C4-C8 (up to 2 rostral segments, intrasegmental, and up to 2 caudal segments), (iii) C7 with stimulation of C4-C8 and L1-L2 (up to 3 rostral segments, intrasegmental, 1 caudal segment, and rostral lumbar), and (iv) C8 with stimulation of C5-C8 and L1-L2 (up to 3 rostral segments, intrasegmental, and rostral lumbar). B. Diagram of the *ex vivo* spinal cord preparation with optogenetic stimulation applied to ipsilateral C4-C8 and L1-L5 whilst recording from sample roots within C4-C8. C. (i) Onset latency, (ii) rate of rise (prior to peak), and (iii) amplitude of motor responses normalized to the maximal response at the recorded root across all stimulation sites recorded from cervical ventral roots with respect to stimulation at rostral (+), intrasegmental (0), and caudal (-) cervical segments (n = 11 mice). Measurements were obtained from averaged traces per stimulation distance and averaged across all animals. 95% CI are shown in error bars. D. (i) Onset latency, (ii) rate of rise (prior to peak), and (iii) amplitude of motor responses normalized to the maximal response at the recorded root across all stimulation sites recorded from cervical ventral roots with respect to stimulation at lumbar segments (<-14 distance) (n = 11 mice). Measurements were obtained from averaged traces per stimulation distance and averaged across all animals. 95% CI are shown in error bars.

Within the cervical cord, we observed that stimulation of local, as well as rostral and caudal segments adjacent to the recorded root, resulted in a uniform, short-latency, high-amplitude, and multi-peaked motor responses (Figure 2A). We measured the onset latency, the rate of rise prior to the peak, and the amplitude of the first major peak scaled to the maximum response recorded at the root across all stimulation sites (Figure 2C). Within a total of 6 adjacent segments (including local), putative motor responses were observed with latencies of about ∼10-12 ms after blue-light stimulation, with peak amplitudes between ∼0.62-0.93 relative to the maximal response, and with a rate of rise between ∼0.035-0.064 V/0.1ms (Figure 2Ci-iii). When recording from C4-C6 ventral roots, putative motor responses were observed when stimulating up to 2 caudal segments (0 to -2 distance) (n = 8 ventral roots, 6 mice; Figure 2Ai: C4 rec, C4-C6 stim; 2Aii: C6 rec, C6-C8 stim). When stimulating past 2 caudal segments (<-2 distance), we observed latencies greater than ∼13 ms, peak amplitudes below ∼0.30 of the maximal response, and a rate of rise below 0.009 V/0.1ms (Figure 2Ci-iii), which we assume are polysynaptic responses based on the longer latencies, lower amplitudes, and the lower rate of rise. On the other hand, when recording from C5-C8 ventral roots, short-latency, fast-rising, high-amplitude motor responses were observed when stimulating up to 3 rostral segments (0 to 3 distance) (n = 16 ventral roots, 12 mice; Figure 2Aiii: C7 rec, C4-C7 stim; 2Aiv: C8 rec, C5-C8 stim). When stimulating past 3 rostral segments (>3 distance), we observed latencies greater than ∼16 ms, peak amplitudes below ∼0.37 of the maximal root response, and a rate of rise below 0.009 V/0.1ms (Figure 2Ci-iii), which we again assume are polysynaptic responses based on the longer latencies, weaker amplitudes, and the lower rate of rise. After stimulating segments beyond these specific rostral and caudal distances relative to the recorded cervical root, our visual observations of distinct changes in the waveform of motor responses (Figure 2A), in combination with the changes observed in the measurements of latency, rate of rise, and relative peak amplitude (Figure 2C), suggests that cervical dI3s form short propriospinal pathways that can monosynaptically activate motoneurons locally, 1-2 segments rostrally, and 1-3 segments caudally.

We also observed that stimulation of L1-L2 segments largely activated C7-C8 segments (-14 to -16 distance) (n = 11 ventral roots, 10 mice; Figure 2Aiii & Aiv). These rostral lumbar stimulations elicited weaker motor responses that had longer latencies relative to local cervical stimulations, specifically with latencies between ∼28-32 ms, peak amplitudes between ∼0.21-0.29 of the maximal response, and a rate of rise of ∼0.010 V/0.1ms (Figure 2Di-iii). While these L1-L2 stimulation C7-C8 motor responses were different from local stimulation, stimulations or recordings beyond this limit (<-16 distance) led to drastic changes in the motor responses. Specifically, we observed latencies greater than ∼38 ms, peak amplitudes below ∼0.15 of the maximal response, and a rate of rise below 0.004 V/0.1ms (Figure 2Di-iii), which we assume are polysynaptic responses based on the longer latencies and lower rate of rise. With increasing distance of lumbar stimulation from cervical roots, our visual observations of changes in motor response in the averaged traces (Figure 2A), and our observation of changes in the latency, rate of rise, and peak amplitude (Figure 2D) suggests that direct ascending pathways formed or activated by lumbar dI3s primarily originate from L1-L2 segments and terminate onto C7-C8 motoneurons.

### Lumbar motoneurons receive input from lumbar and cervical dI3s

We then characterized the dI3s that provide input to the lumbar enlargement by independently activating ipsilateral C4-C8 and L1-L5 segments while recording from L1-L5 ventral roots (Figure 3B). We performed the same averaging of short pulse recordings and identified dI3 subpopulations located in the cervical and lumbar segments that form distinct connections with lumbar motoneurons (Figure 3A).

**Figure 3.**
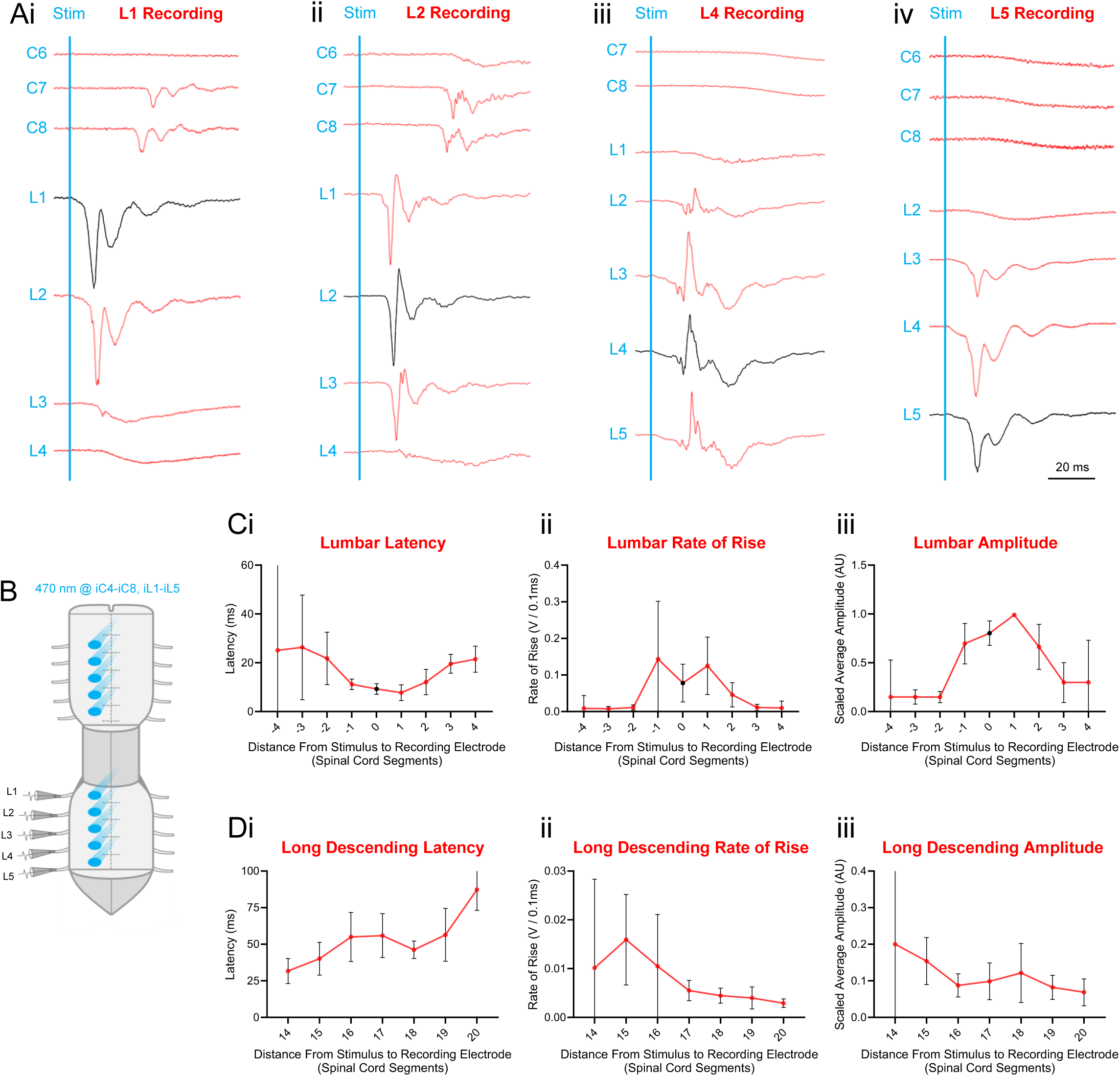
Lumbar motoneurons receive input from lumbar and cervical dI3 neurons. A. Averaged traces of lumbar ventral root recordings in response to 10 ms optogenetic stimulations of ipsilateral dI3 neurons located from segments L1-L5 and C4-C8. Recordings of optogenetic stimulation of intrasegmental dI3s (black trace), rostral dI3s (red traces above), and caudal dI3s (red traces below) relative to the lumbar ventral root are shown. Short-latency, fast-rising, high-amplitude motor responses were observed in: (i) L1 with stimulation of C7-C8 and L1-L2 (caudal cervical, intrasegmental, and 1 caudal segment), (ii) L2 with stimulation of C7-C8 and L1-L3 (caudal cervical, 1 rostral segment, intrasegmental, and 1 caudal segment), (iii) L4 with stimulation of L2-L5 (up to 2 rostral segments, intrasegmental, and 1 caudal segment), and (iv) L5 with stimulation of L3-L5 (up to 2 rostral segments and intrasegmental). B. Diagram of the *ex vivo* spinal cord preparation with optogenetic stimulation applied to ipsilateral C4-C8 and L1-L5 whilst recording from sample roots within L1-L5. C. (i) Onset latency, (ii) rate of rise (prior to peak), and (iii) amplitude of motor responses normalized to the maximal response at the recorded root across all stimulation sites recorded from lumbar ventral roots with respect to stimulation at rostral (+), intrasegmental (0), and caudal (-) lumbar segments (n = 11 mice). Measurements were obtained from averaged traces per stimulation distance and averaged across all animals. 95% CI are shown in error bars. D. (i) Onset latency, (ii) rate of rise (prior to peak), and (iii) amplitude of motor responses normalized to the maximal response at the recorded root across all stimulation sites recorded from lumbar ventral roots with respect to stimulation at cervical segments (>14 distance) (n = 11 mice). Measurements were obtained from averaged traces per stimulation distance and averaged across all animals. 95% CI are shown in error bars.

Similar to the cervical segments, within the lumbar enlargement we observed uniform, short-latency, fast-rising, high-amplitude, and multi-peaked motor responses with stimulations of local, rostral, and caudal segments adjacent to specific ventral roots (Figure 3A). We measured the onset latency, the rate of rise prior to the peak, and the amplitude of the first major peak scaled to the maximum response recorded at the root across all stimulation sites (Figure 3C). We observed putative monosynaptic motor responses within a total of 4 adjacent segments (including local), ∼8-12 ms after the blue-light stimulation, with peak amplitudes between ∼0.66-0.99 relative to the maximal response, and a rate of rise between ∼0.046-0.143 V/0.1ms (Figure 3Ci-iii). When recording from L1-L4 roots, we observed these short-latency, fast-rising, high-amplitude motor responses with stimulation of dI3s up to only 1 caudal segment (0 to -1 distance) (n = 12 ventral roots, 11 mice; Figure 3Ai: L1 rec, L1-L2 stim; 3Aii: L2 rec, L2-L3 stim). When stimulating more caudally (<-1 distance), we observed latencies greater than ∼22 ms, peak amplitudes lower than ∼0.15 of the maximal response at the root across all simulations sites, and a rate of rise below 0.011 V/0.1ms (Figure 3Ci-iii). Conversely, short-latency, fast-rising, high-amplitude motor responses could be elicited in more caudal L2-L5 roots with stimulations of dI3s up to 2 rostral segments away (0 to 2 distance) (n = 14 ventral roots, 11 mice; Figure 3Aiii: L4 rec, L2-L4 stim; 3Aiv: L5 rec, L3-L5 stim). When stimulating more rostrally (>2 distance), we observed latencies greater than ∼20 ms, peak amplitudes lower than ∼0.30 of the maximal response, and a rate of rise below 0.011 V/0.1ms (Figure 3Ci-iii), which we assume are polysynaptic responses based on the longer latencies and lower rate of rise. Our observations of changes in motor responses in the averaged traces (Figure 3A) when stimulating segments beyond a specific rostral and caudal distance relative to the recorded lumbar root, in combination with the changes in latencies, peak amplitude, and rate of rise (Figure 3C) suggests that within the lumbar enlargement, short propriospinal dI3s can monosynaptically activate motoneurons intrasegmentally, 1 segment rostrally, and 1-2 segments caudally.

Next, we observed that stimulation of C7-C8 segments largely activated L1-L2 segments (14-16 distance) (n = 11 ventral roots, 10 mice; Figure 3Ai & Aii). While these cervical simulations elicited longer latency and weaker motor responses relative to local stimulation in the lumbar enlargement, specifically with latencies between ∼32-40 ms, peak amplitudes between ∼0.15-0.20 of the maximal response at the root across all simulations sites, and a rate of rise of ∼0.010-0.016 V/0.1ms (Figure 3Di-iii), these motor responses were markedly distinct from longer distance stimulations relative to the recorded root based on latency, rate of rise, and amplitude. When stimulating more rostrally in the cervical enlargement or recording more caudally in the lumbar (>16 distance), motor responses had latencies greater than ∼55ms, peak amplitudes lower than ∼0.12 of the maximal response at the root across all simulations sites, and a rate of rise below ∼0.006 V/0.1ms (Figure 3Di-iii). Based on our observations in the averaged traces (Figure 3A), in combination with the latencies, rate of rise, and amplitude measured across these distances (Figure 3D), we suggest that long descending propriospinal pathways formed or activated by dI3s originate from C7-C8 and terminate onto L1-L2 segments. Furthermore, our observations of both long ascending and descending connectivity patterns suggest that dI3s reciprocally activate C7-C8 and L1-L2 segments.

### dI3 neurons activate contralateral motoneurons

To the best of our knowledge, dI3s do not form any commissural projections (Avraham et al., 2010; Goetz et al., 2015). However, we observed strong motor responses when unilaterally stimulating contralateral cervical and lumbar segments (Figure 4B). These contralateral stimulations visually produced the same pattern of activity across the spinal cord as seen in the ipsilateral stimulations in the averaged recordings (Figure 4A). We measured the same parameters as previously to characterize the differences between the ipsilateral and contralateral motor responses across the spinal cord. The contralateral motor responses had a slightly longer latency (n=11 mice, 0.7907 ms, p<0.001, 95%CI=0.4188 to 1.163), slightly weaker peak amplitude (n=11 mice, -0.03534 relative to maximal response, p<0.001, 95%CI=-0.04974 to -0.02094), and a slightly smaller rate of rise (p=11 mice, - 0.03534 V/0.1ms, p<0.001, 95%CI=-0.04974 to -0.02094) (Figure 4Ci-iii). While there were small differences within the same segment, we observed the same monosynaptic and polysynaptic activation pattern across the spinal cord. Specifically, within the cervical cord, the short monosynaptic responses were observed with stimulations up to 3 rostral segments away and up to 2 caudal segments away (data not shown), while within the lumbar cord, dI3 subpopulations activated contralateral motoneurons from up to 2 rostral segments away and up to 1 caudal segment away (data not shown). Finally, we also observed the same cervicolumbar and lumbocervical activation patterns on the contralateral side, where C7-C8 dI3s activated contralateral L1-L2 segments and L1-L2 dI3s activated contralateral C7-C8 segments (Figure 4A).

**Figure 4.**
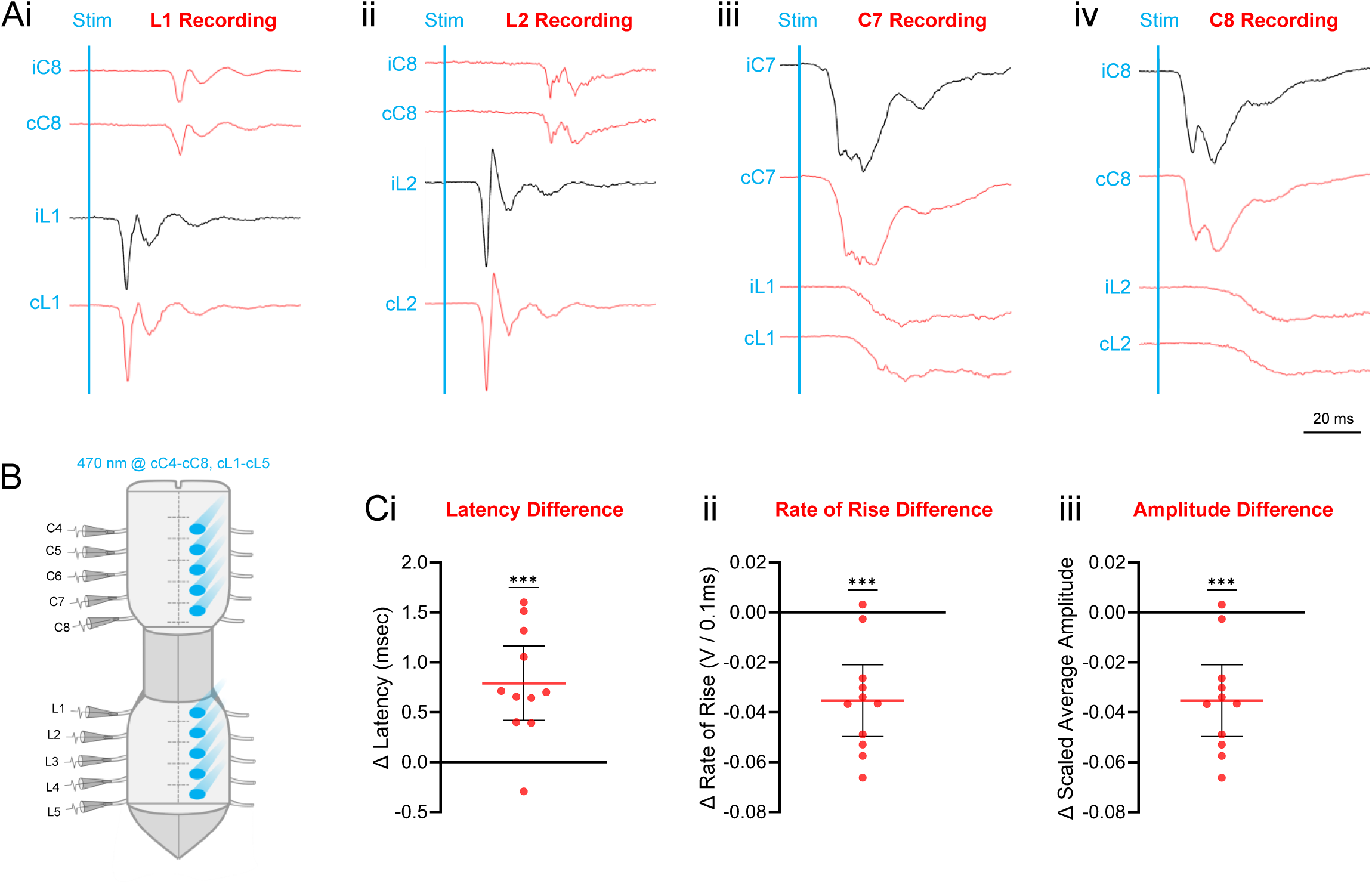
dI3 neurons activate contralateral motoneurons. A. Averaged traces of lumbar and cervical ventral root recordings in response to 10 ms optogenetic stimulations of ipsilateral and contralateral dI3 neurons located from segments L1-L5 and C4-C8. Recordings of optogenetic stimulation of ipsilateral intrasegmental (black), ipsilateral non-intrasegmental (red), and contralateral (red) relative to the ventral root recording are shown. Stimulation of dI3 neurons located within the same segment contralaterally or ipsilaterally elicits the same motor response across the spinal cord. Short-latency, fast-rising, and high-amplitude motor response was observed in: (i) L1 with stimulation of C8 and L1 ipsilateral and contralateral pairs, (ii) L2 with stimulation of C8 and L2 ipsilateral and contralateral pairs, (iii) C7 with stimulation of C7 and L1 ipsilateral and contralateral pairs, and (iv) C8 with stimulation of C8 and L2 ipsilateral and contralateral pairs. B. Diagram of the *ex vivo* spinal cord preparation with optogenetic stimulation was applied to contralateral C4-C8 and L1-L5 whilst recording from sample roots within L1-L5. C. The difference in onset latency (i), rate of rise (prior to peak) (ii), and amplitude (iii) between motor responses from ipsilateral and contralateral stimulation of dI3s. Difference is calculated as following: contralateral-ipsilateral. Significant differences from 0 indicated with ***, p<0.001, one sample t-test. 95% CI are shown.

### dI3 neurons activate lumbar locomotor networks

Bui et al., 2016 provided indirect evidence that dI3s provide excitatory inputs to locomotor networks in the lumbar spinal cord. They recruited cutaneous afferents via sural nerve stimulations in control and dI3-silenced spinal cord preparations and observed that the dI3-silenced spinal cords had deficits in producing locomotor activity in the ventral roots (Bui et al., 2016). To determine which dI3 subpopulations are directly involved in activating locomotor networks, we applied 10-second-long blue-light stimulations at individual unilateral segments of the lumbar cord and recorded from rostral (L1-L2) and caudal (L4-L5) ventral roots, generally associated with flexors and extensors, respectively (Toossi et al., 2019). We observed that dI3 subpopulations located throughout L1-L5 can independently recruit locomotor networks and produce alternating, rhythmic activity between flexor and extensor roots in the lumbar cord (n = 5/5 mice; Figure 5A).

**Figure 5.**
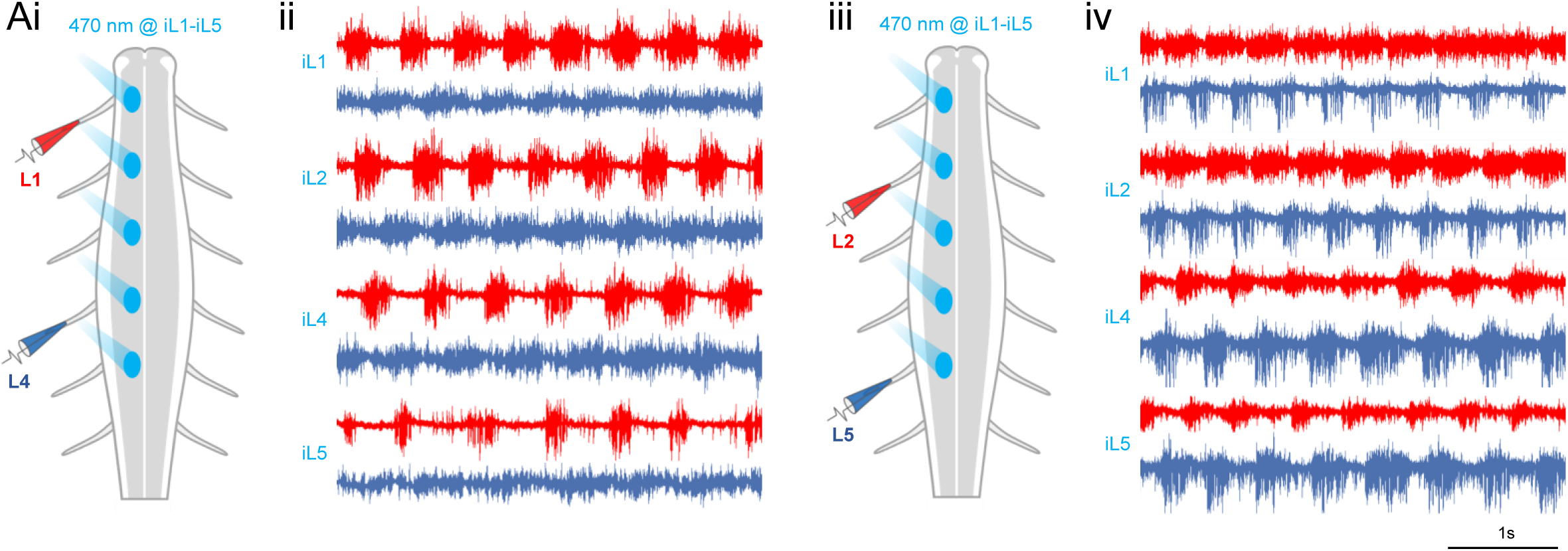
dI3 neurons activate lumbar locomotor networks. A. Optogenetic stimulation of ipsilateral lumbar dI3s evokes fictive locomotion in flexor and extensor roots. (i) and (iii) Experimental setup and sample locomotor recordings. Spinal cord preparations consisted of isolated lumbar spinal cords from neonatal dI3-driver-CatCh mice with a dorsal hemisection. Suction electrodes were applied to (i) L1 and L4 (red) and (iii) L2 and L5 (blue) roots while optogenetically stimulating ipsilateral L1-L5 segments (L3 not shown). (ii and iv) Blue light was applied for 10 seconds continuously, and alternating, rhythmic activity was observed in both recording electrodes during the stimulation period.

## Discussion

dI3 neurons are found throughout the cervical and lumbar spinal cord, and their connectivity to motoneurons is well established (Bui et al., 2013; Goltash et al., 2023; Stepien et al., 2010); however, how dI3s connect across the span of their respective cervical or lumbar enlargement and between these two major divisions of the nervous system responsible for the control of forelimb and hindlimb motor function, respectively (Pocratsky et al., 2020; Ruder et al., 2016; Shepard et al., 2023), is not well characterized. In this study, we used optogenetic expression of CatCh within dI3 neurons to perform circuit mapping experiments in *ex vivo* neonatal spinal cord preparations. Patterns and amplitudes of motor responses within the cervical or lumbar spinal cord or across these regions to dI3 stimulation may provide insights into how dI3s shape forelimb and hindlimb motor function through segmental, inter-segmental, and inter-enlargement connections.

### Cervical dI3 neurons form short propriospinal circuits for forelimb function

Forelimb motor control is partly facilitated by cervical propriospinal neuron populations that integrate various supraspinal and sensory commands and activate motor pools across the cervical cord (Alstermark & Isa, 2012; Azim et al., 2014; Bui et al., 2013; Ueno et al., 2018). We found that cervical dI3 subpopulations activate a widespread number of segments. Specifically, we found that a single dI3 subpopulation within a cervical segment can project to a total of 6 adjacent cervical segments, with a slight descending bias (Figure 6Ai). We also observed multiple peaks in the motor responses and weak responses in more distal segments, suggesting that dI3s also activate polysynaptic pathways within the cervical enlargement (Figure 6Ai).

**Figure 6.**
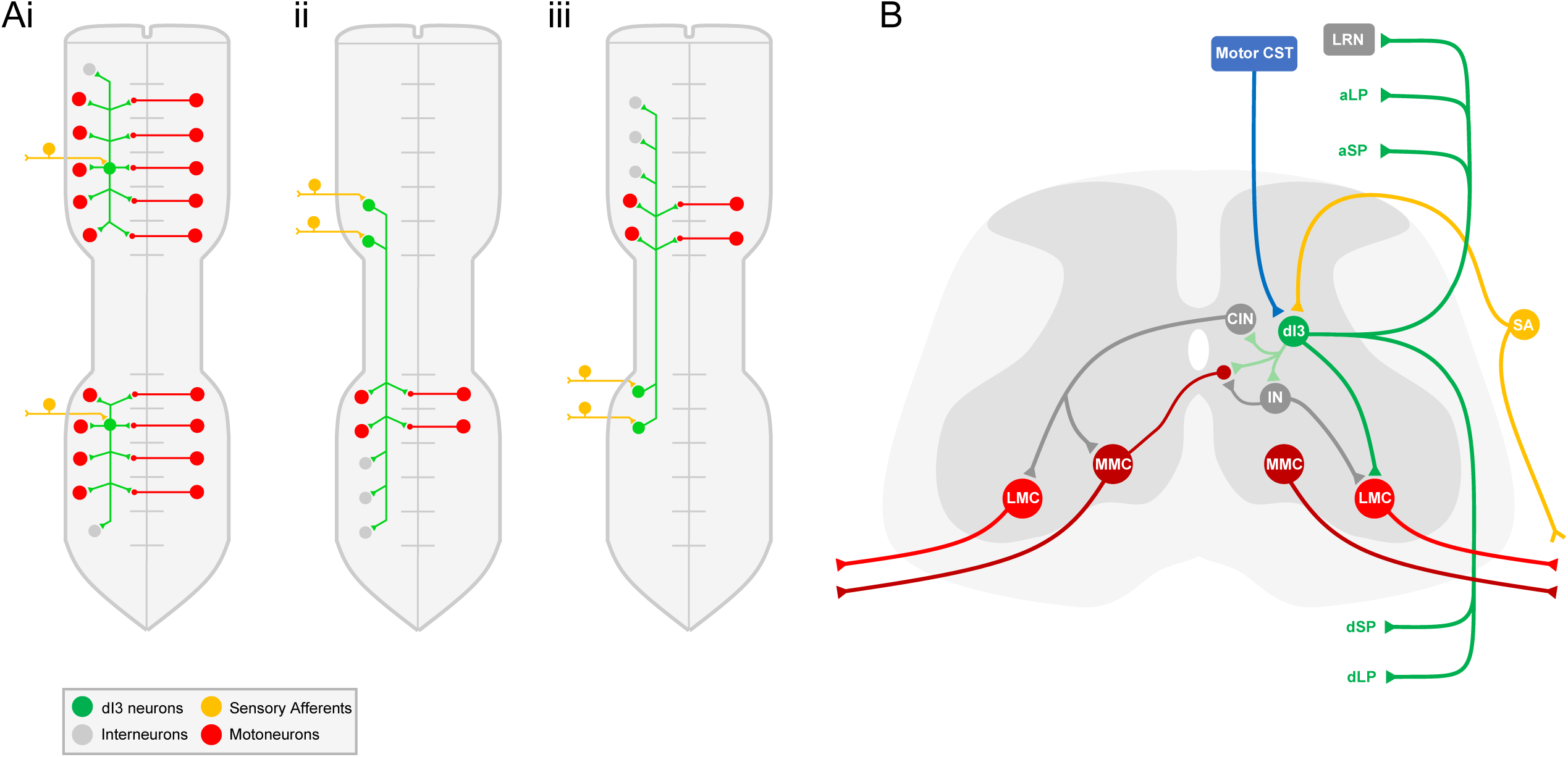
dI3 neurons form sensorimotor circuits across the spinal cord for motor control. A. Summary diagram of rosto-caudal connectivity patterns of dI3 neurons in the cervical and lumbar enlargments. (i) Cervical dI3s can project intrasegmentally, up to 2 segments rostrally, up to 3 segments caudally to motoneurons, and polysynaptically activate more distal segments. Lumbar dI3s can project intrasegmentally, 1 segment rostrally, up to 2 segments caudally to motoneurons, and polysynaptically activate more distal segments. (ii) Cervical dI3s in C7-C8 form long descending propriospinal pathways to L1-L2 motoneurons and polysynaptically activate more caudal lumbar segments polysynaptically. (iii) Lumbar dI3s in L1-L2 form long ascending propriospinal pathways to C7-C8 motoneurons and activate more rostral cervical segments polysynaptically. B. Summary of dI3 connectivity patterns across the spinal cord to various networks for motor control. dI3 neurons receive inputs from cutaneous mechanoreceptive and proprioceptive sensory afferents (SA) (Bui et al., 2013) and motor corticospinal tracts (Motor CST) from the primary motor cortex (Ueno et al., 2018). dI3s directly activate intrasegmental motoneurons primarily in the lateral motor column (LMC) and activate ipsilaterally projecting interneurons (IN) that activate the LMC (Bui et al., 2013; Goetz et al., 2015; Stepien et al., 2010). dI3s activate motoneurons beyond their segment through ascending and descending short propriospinal (aSP and dSP) and long propriospinal (aLP and dLP) projections. dI3s relay sensory commands to the lateral reticular nucleus (LRN) via long ascending projections (Pivetta et al., 2014). Proposed circuitry of dI3s activating contralateral segments: dI3s directly activate commissural dendrites of contralateral motoneurons in the medial motor column (MMC), activate ipsilaterally projecting interneurons (IN) that activate commissural dendrites of contralateral MMC motoneurons, and activate commissural interneurons (CIN) that activate the contralateral MMC and LMC (Goetz et al., 2015).

The dI3 neurons are known to be critical for the control of hand grasp (Bui et al. 2013); however, since dI3 subpopulations likely integrate proprioceptive and cutaneous sensory feedback from different muscles and areas of the skin, their widespread innervation patterns suggest that they may also shape, in an accessory manner, a wide range of forelimb movements such as reaching and forelimb locomotion. Additionally, a major excitatory propriospinal neural population within the cervical cord are V2a’s, which have been found to facilitate skilled reaching behaviour (Azim et al., 2014). While inhibition of dI3s did not lead to marked deficits in skilled reaching tasks (Bui et al., 2013), our observations of multiple peaks during ipsilateral motor responses suggests that they activate ipsilaterally projecting interneurons. Through this polysynaptic activation of potentially other spinal populations, such as V2as, dI3s may interact with other networks to incorporate grasping function with skilled reaching to coordinate complete movements.

### Lumbar dI3 neurons form short propriospinal circuits for hindlimb function

Similarly, the lumbar propriospinal circuitry forms specific networks that integrate sensory feedback to modulate different networks to maintain balance and coordination in different contexts (Britz et al., 2015; Bui et al., 2016; Danner et al., 2019; Dyck et al., 2012; Goetz et al., 2015; Koch et al., 2017; Lanuza et al., 2004; Talpalar et al., 2013; H. Zhang et al., 2022; J. Zhang et al., 2014; Y. Zhang et al., 2008); however, we still have a limited understanding of how these inputs are integrated by propriospinal neurons to modulate the activity of specific motor pools to shape motor output. We found that lumbar dI3 subpopulations can directly project up to a total of 4 adjacent segments and identified a descending bias in their connectivity (Figure 6Ai). Specifically, a dI3 subpopulation could activate motoneurons located 1 rostral segment away and motoneurons located up to 2 segments caudally (Figure 6Ai). We also observed that dI3s can activate polysynaptic networks locally and beyond their monosynaptic activation range, suggesting that they can also interact with other locomotor or propriospinal networks for different functions (Figure 6Ai).

Given the general flexor-to-extensor gradient in the rostral-to-caudal direction within the lumbar cord (Toossi et al., 2019), this descending bias in dI3 projections suggests that dI3s may be involved in facilitating swing-to-stance transitions during locomotion. Silencing of dI3s *in vivo* led to deficits in hindlimb placement during horizontal ladder walking (Bui et al., 2013), and inhibition of dI3s during locomotion altered the stance and swing phases indicating a disruption in the timing of flexor and extensor muscles (Bui et al., 2016). Additionally, these mice had a severe loss in weight support, marked by an increase in hindlimb foot spacing, and after a complete thoracic spinal cord injury, dI3-silenced mice lost any ability in promoting functional recovery (Bui et al., 2016). It is likely that weight support and the step transitions facilitated by the multi-segmental and descending connectivity of lumbar dI3s play an essential role in promoting recovery.

### Cervical dI3 neurons form long descending propriospinal circuits

Long descending PINs are localized throughout the cervical and thoracic spinal cord (Ni et al., 2014). Within the cervical enlargement, commissural and ipsilaterally projecting long descending neurons are localized throughout C1-C8 and project to the rostral lumbar cord (L1-L3) (Brockett et al., 2013; Reed et al., 2006). Depending on their localization, sources of proprioceptive and cutaneous inputs, and axonal projection patterns, these long descending PINs function to fine-tune locomotion, stabilize posture, and maintain interlimb coordination by activating specific hindlimb muscles based on neck, trunk, and forelimb sensory feedback (Mayer & Akay, 2021; Ruder et al., 2016; Shepard et al., 2023). Silencing these long descending pathways led to significant deficits in forelimb-hindlimb coordination and left-right coordination in both forelimb and hindlimb pairs; however, these deficits in left-right coordination were more apparent in the hindlimbs (Shepard et al., 2023).

We identified a subpopulation of dI3s within the C7-C8 segments that distinctly activate L1-L2 segments of the lumbar cord; however, we also observed polysynaptic motor responses in L1-L2 segments with the stimulation of C4-C6 dI3s and observed similar polysynaptic activity in more caudal lumbar segments (L3-L5) with the stimulation of C4-C8 dI3s (Figure 6Aii). Ruder et al., 2016 demonstrated that there were a limited number of dI3s throughout C4-C7 with anatomical projections to the lumbar segments; however, the strong response we observed at L1-L2 with C7-C8 stimulation could potentially involve a small subpopulation of dI3s or a distinct dI3-activated pathway involving other long descending PIN populations, such as V2a and Shox2 neurons (Flynn et al., 2017; Ni et al., 2014; Ruder et al., 2016). Furthermore, the polysynaptic recruitment of other long descending networks from the stimulation of dI3s across the cervical enlargement suggests that cervical dI3s can influence a wide range of movements in the hindlimbs. Similar to Shepard et al., 2023, where hindlimb coordination was hindered with the inhibition of long descending pathways, silencing of dI3s led to distinct deficits in hindlimb placement during horizontal ladder tasks (Bui et al., 2013). In combination with lumbar dI3 sensorimotor circuits, these dI3-mediated long descending pathways may integrate supraspinal and forelimb sensory inputs (Bui et al., 2013; Mayer & Akay, 2021; Ueno et al., 2018) to facilitate proper hindlimb movement.

### Lumbar dI3 neurons form long ascending propriospinal circuits

Long ascending PINs are primarily located in the L1-L3 segments and can project to C5-C8 segments (Reed et al., 2006); however, a majority of these projections terminate in the ventrolateral motor nuclei of C7-C8 that supplies the axilla muscles (Brockett et al., 2013). The functional role of long ascending propriospinal pathways is primarily for interlimb coordination during locomotion (Pocratsky et al., 2020). Inhibition of long ascending propriospinal pathways led to deficits in forelimb-hindlimb coordination and disrupted left-right coordination in both forelimb and hindlimb pairs; however, these deficits were highly context-dependent, specifically occurring with enhanced sensory input via a high friction surface during treadmill locomotion (Pocratsky et al., 2020).

While other excitatory PIN subclasses form long ascending pathways, such as Shox2, V3, and V0_V_ INs (Ruder et al., 2016; H. Zhang et al., 2022), these populations likely receive different sources of sensory input and have slightly different projection characteristics (Laliberte et al., 2019). While it is not currently clear whether dI3s can form these long ascending pathways independently, our observations of strong activity at C7-C8 segments with stimulation of L1-L2 dI3s suggest that they directly form or contribute to the activation of these pathways (Figure 6Aiii). Furthermore, we saw strong recruitment of polysynaptic networks at more rostral cervical segments (C4-C6) with stimulation of lumbar dI3s (L1-L5) and at C7-C8 with stimulation of more caudal lumbar (L3-L5) dI3s. These observations suggest that dI3s likely relay hindlimb sensory inputs to influence the activity of other long ascending networks that coordinate different forelimb-hindlimb muscles.

The combined role of the long ascending and descending pathways may be involved in forming a loop between the cervical and lumbar enlargements to coordinate left-right alternations in the hindlimb and forelimb pairs (Shepard et al. 2023). Our findings of distinct reciprocal ascending and descending pathways formed by dI3s between L1-L2 and C7-C8 suggest that they play a role in coordinating a distinct set of muscle groups between the 2 enlargements. It is notable that inhibition of dI3s had a significant effect on the homolateral interlimb coordination during treadmill locomotion (Bui et al. 2016).

### dI3 neurons activate commissural pathways for postural stability

While studies have shown that dI3s do not directly form commissural projections, it was surprising to observe short latency, high-amplitude motor responses with unilateral stimulations of contralateral spinal segments. We observed a minor difference in latency of just under ∼1 ms in these contralateral responses, corresponding to potentially a single interposed synapse or increased conduction time with the longer commissural pathway. Since the rostrocaudal activation pattern was identical on both sides, there are multiple possibilities regarding the indirect activation of these specific contralateral motoneurons.

Contralateral motor responses could result from direct optogenetic stimulation of contralateral dI3s with crossing commissural dendrites expressing CatCh or it can be due to synaptic excitation of crossing commissural dendrites of contralateral motoneurons by stimulated ipsilateral dI3s. Since *Isl1* is expressed in both dI3s and motoneurons (Bui et al., 2013), experiments performing unilateral intraspinal injections of AAV-FRT-FP into *Isl1^Cre^*; *Tau^FLP^* mice were able to visualize dendritic patterns of both dI3s and motoneurons (Goetz et al., 2015). The dI3 neurons do not seem to have any significant commissural dendrites; however, motoneurons in the medial motor column (MMC) have a significant number of dendrites that innervate the contralateral side (Goetz et al., 2015). Furthermore, these commissural MN dendrites also received synaptic input from interneurons that exclusively project ipsilaterally, such as V2 and V1s (Goetz et al., 2015). This provides further support that dI3s likely innervate these commissural MN dendrites to facilitate a short latency motor response in the same contralateral segments that are activated on the ipsilateral side (Figure 6B).

This symmetrical activation of contralateral segments likely has functional implications of coordinating left and right networks for different purposes. While ipsilaterally projecting INs, including dI3s, primarily activate the ipsilateral lateral motor column (LMC), which is involved in controlling various limb muscles, they innervate the commissural dendrites of the contralateral MMC, which is involved in modulating axial muscles for postural stability (Goetz et al., 2015). Conversely, excitatory commissural INs have access to the contralateral MMC and LMC (Goetz et al., 2015) and play a role in synchronizing left and right limb movements for highspeed locomotion. For example, commissural IN populations, such as the V3 and V0_V_ INs, function to promote left-right coordination during high-speed locomotion by increasing the synchronization of ipsilateral and contralateral limb muscles (Danner et al., 2019; Talpalar et al., 2013; H. Zhang et al., 2022). While the role of these dI3-activated commissural pathways has not been tested, they likely play a role in activating contralateral axial muscles to facilitate postural stability and may not be directly involved in crossed reflexes or left-right limb coordination. However, since the commissural INs that activate the contralateral LMC do not independently activate ipsilateral LMCs (Goetz et al., 2015), dI3s may be needed to synchronize the activation of the ipsilateral LMC with these commissural INs during high-speed locomotion (Figure 6B).

### dI3s interneurons activate rhythm generating circuits

Our findings suggest that dI3s localized across the lumbar cord can recruit locomotor activity with tonic blue light stimulation. The rhythm-generating network highly involves Shox2 neurons (Dougherty et al. 2013). While the majority of Shox2 neurons are Chx10-expressing V2a neurons, ablation of Shox2 V2as had minimal effects on the locomotor rhythm and mainly led to deficits in burst amplitude and burst duration; however, a subset of non-V2a Shox2 neurons that do not express Chx10, which includes a small population of Isl1-positive dI3s, are involved in producing locomotor rhythm (Dougherty et al. 2013). Together, the non-V2a Shox2 neurons provide rhythmic drive to Shox2 V2a neurons, which modulate burst amplitude and duration (Dougherty et al., 2013; Kiehn, 2016). From our observations, it appears that dI3s provide excitatory drive to non-V2a Shox2 neurons to generate locomotor rhythm.

Shox2 neurons have a strong flexor bias in terms of their connectivity and their rostral populations are rhythmically active during locomotion (Dougherty et al. 2013). Although these neurons are electrically coupled via gap junctions in neonates (Ha et al. 2018), descending drive of locomotor activity likely requires other IN populations with intersegmental projections. Since dI3s are rhythmically active during locomotion (Bui et al. 2016), the descending bias in their monosynaptic and polysynaptic connectivity may contribute to propagating this locomotor rhythmic activity towards caudal segments, further supporting their involvement in facilitating transitions from flexors to extensors, as previously described.

## Methods

### Animals

All animal protocols and experimental procedures were followed with respect to the guidelines of the Canadian Council of Animal Care (CCAC) and have been approved by the University of Ottawa Animal Care Committee (BL-3945). The dI3 neurons were targeted in mice through Cre and FlpO recombinase expression driven by *Isl1* and *Slc17a6* (Vglut2), respectively. *Isl1^Cre^*^+/-^ mice were crossed with *Slc17a6^FlpO^*^+/+^ mice to drive the expression of Cre and FlpO recombinase in dI3 neurons and Vglut2^+^ nociceptive afferents, forming dI3-driver mice. For optogenetic experiments, *Isl1^Cre^*^+/-^; *Slc17a6^FlpO^*^+/+^ (dI3-driver) mice were crossed with *Ai80^(RCFL-CatCh)-D^* (CatCh) mice (The Jackson Laboratory, Strain No.: 025109) containing a *frt*-flanked STOP cassette and a *loxP*-flanked STOP cassette upstream of the CatCh/EYFP fluorescent protein, resulting in *Isl1^Cre^*^+/-^; *Slc17a6^FlpO^*^+/+^; *Ai80^(RCFL-CatCh)-D^* (dI3-driver-CatCh) mice. To visualize the dI3 neurons and Vglut2^+^ afferents, dI3-driver mice were crossed with *Ai65^(RCFL-tdT)-D^* (TdTomato) mice (The Jackson Laboratory, Strain No.: 021875) containing a *frt*-flanked STOP cassette and a *loxP*-flanked STOP cassette upstream of the TdTomato fluorescent protein, resulting in *Isl1^Cre^*^+/-^; *Slc17a6^FlpO^*^+/+^; *Ai65^(RCFL-tdT)-D^* (dI3-driver-TdTomato) mice.

### Spinal Cord Preparation

Postnatal (P0-P7) mice were anesthetized using euthanyl (Bimeda-MTC; 1.85 mL/kg for mice at a dilution of 120 mg/mL) via an intraperitoneal injection followed by decapitation. Complete spinal cords were isolated via a laminectomy in oxygenated and room temperature recording artificial cerebrospinal fluid (aCSF) and dissected by cutting the dorsal and ventral roots as distally as possible. Recording aCSF (in mM: NaCl, 127; KCl, 3; NaH2PO4, 1.25; MgCl2, 1; CaCl2, 2; NaHCO3, 26; D-glucose, 10) was made daily using diH2O, oxygenated at room temperature, and pH and osmolarity tested. For experiments requiring the removal of CatCh^+^ nociceptive afferents, a dorsal hemisection was performed at lumbar and cervical enlargements by pinning the spinal cord on its side and making rostrocaudal incisions using surgical blades approximately one-third of the way along the dorsoventral axis. For locomotor recordings, the lumbar spinal cord was isolated. Spinal cords were rested for at least 1 hour in oxygenating recording aCSF before getting pinned to a base of clear Sylgard (Dow Corning) in a recording chamber while perfused with fresh circulating oxygenated aCSF at room temperature. A Minipuls 3 peristaltic pump (Gilson; F155005) was used to circulate ACSF at a flow rate of over 3 mL/min.

### Optogenetics

Stimulation of dI3 neurons was performed using the *Isl1^Cre^*^+/-^; *Slc17a6^FlpO^*^+/+^; *Ai80^(RCFL-CatCh)-D^* (dI3-driver-CatCh) mice and blue light (X-Cite 120 LEDMini; 470 nm). In the dorsal hemisection preparation, dI3 subpopulations were stimulated by focusing the light on approximately one unilateral spinal segment from C4-C8 and L1-L5 of the ipsilateral and contralateral sides of the spinal cord relative to the roots being recorded. Since CatCh causes rapid depolarization, a 10 ms pulse protocol was used for recordings of short-latency connectivity to motoneurons to avoid activation of locomotor networks, and for locomotor recordings, a 10-second pulse protocol was used at 50% light intensity.

### Extracellular Recordings

For ventral root recordings, 1.50 mm/1.10 mm (OD/ID) borosilicate glass capillary tubes (Sutter Instruments; BF150-110-10) were pulled to a selected length using a P-2000 micropipette puller (Sutter Instruments) and the tips were broken to fit the ventral roots. The micropipette tips were then attached to bipolar suction electrodes (A-M Systems; 573040) and filled to the electrode tip with aCSF before suctioning the roots. Ventral roots from C4-C8 and L1-L5 were recorded for cervical and lumbar segments. Recordings were amplified 10,000X in differential mode and bandpass filtered from 1 Hz to 3 KHz (A-M Systems; Model 1700 Differential AC Amplifier; 690000) and acquired at 10kHz (Molecular Devices; pClamp nine software, RRID:SCR_011323; Digidata 1550 Data Acquisition System, DD1550).

### Data Analysis

Signal filters were applied to short pulse recordings to remove electrical and high frequency noise. The electrical interference filter removed 60Hz with a 4Hz 3dB bandwidth up to 20 harmonics, and to remove high frequency noise, a Gaussian low-pass filter at 1000Hz was applied. Each short pulse recording consisted of 10 sweeps; after filtering, these sweeps were averaged to produce an averaged trace, that was then rectified and time-integrated. From the processed traces, we obtained the latency of the onset and peak of motor response to calculate the rate of rise. Additionally, we measured the amplitude of the highest peak and normalized all the recordings from the same ventral root to the highest amplitude response for that root across all stimulation sites. Projection distance was obtained by subtracting the stimulation segment from the recording segment, with positive projection distances denoting rostral stimulation sites and negative projection distances denoting caudal stimulation sites.

### Statistics

The statistical analysis was performed in Prism7 (GraphPad Software, Inc.). One sample t-tests were used to compare differences in latency, rate of rise, and amplitude between ipsilateral and contralateral stimulations.

## Acknowledgements

This research was funded by a Canadian Institute of Health Research Project Grant (PJT 180556) and a Natural Sciences and Engineering Research Council Discovery Grant (NSERC RGPIN-2022-03898).

## Contributions

S.N., A.M.L., and T.V.B. conceptualized the paper; S.N. designed and performed ex vivo spinal cord optogenetics and ventral root recordings; A.M.L. analyzed the data and performed all statistics; S.N. wrote the paper and designed the figures, with editing from T.V.B.; T.V.B. supervised this work as a senior author.

